# Transcranial Brain Atlas Based on Photon Measurement Density Function in a Triple-Parameter Standard Channel Space

**DOI:** 10.1101/2024.06.06.597588

**Authors:** Lijiang Wei, Yang Zhao, Farui Liu, Yuanyuan Chen, Yilong Xu, Zheng Li, Chaozhe Zhu

## Abstract

Functional near-infrared spectroscopy (fNIRS) is a widely used transcranial brain imaging technique in neuroscience research. Nevertheless, its lack of anatomical information poses challenges for designing appropriate optode montage and localizing fNIRS signals to underlying anatomical regions. The photon measurement density function (PMDF) is often employed to address these issues, as it accurately measures the sensitivity of a fNIRS signal channel to perturbations of absorption coefficients at any brain locations. However, existing PMDF-based localization methods have two limitations: a limited channel space, and PMDF estimation based on single standard head models, which differ anatomically from individual subjects. To overcome these limitations, this study proposes a continuous fNIRS standard channel space and constructs a PMDF-based transcranial brain atlas (PMDF-TBA) by calculating PMDFs based on MRI images of 48 adults. PMDF-TBA contains group-averaged sensitivities of channels to gray matter and brain regions of atlases, such as Brodmann, AAL2, and LPBA40. Through leave-one-out cross-validation, we evaluated the prediction ability of PMDF-TBA for sensitivity of unseen individuals and found that it outperformed PMDFs based on single standard head models. Thus, PMDF-TBA serves as a more generalizable fNIRS spatial localization tool. Therefore, PMDF-TBA can be utilized to optimize optode montage design, improve channel sensitivity to target brain regions, and assist in the source localization of fNIRS data, thereby promoting the application of fNIRS in neuroscience research.

## 1. Introduction

Functional near-infrared spectroscopy (fNIRS) is a transcranial brain imaging technique that indirectly reflects neural activity by emitting near-infrared light through the scalp and skull to detect changes in cortical blood oxygenation levels (Ferrari and Quaresima, 2012). fNIRS devices are lightweight, portable, low-cost and ecologically valid, making them suitable for special populations such as infants (Collins-Jones et al., 2021), the elderly (Mason et al., 2022), and patients with neuropsychiatric disorders (Obrig, 2014). Due to these advantages, fNIRS has been widely applied in basic and clinical neuroscience research in recent years, and is becoming one of the essential tools for brain function imaging (Pinti et al., 2020).

fNIRS imaging technology lacks inherent anatomical information, posing significant challenges in practical applications. These challenges can be categorized into two main aspects: (1) designing appropriate optode montage prior to experiments to accurately measure target brain regions (Brigadoi et al., 2018; Fu and Richards, 2021; Zimeo Morais et al., 2018), and (2) localizing the anatomical brain regions corresponding to the activated fNIRS channels post-experiment to facilitate meaningful interpretation of the results (Aasted et al., 2015; Cai et al., 2021). An increasing number of researchers have begun to focus on these issues and seek solutions.

The balloon inflation model has been the traditional method for localizing the anatomical brain regions corresponding to fNIRS channels. This model assumes that the cortical surface point directly beneath the midpoint of the line connecting the source and detector optodes is the primary source of the channel signal (Okamoto and Dan, 2005). Researchers further refined and improved this model (Tsuzuki and Dan, 2014). More recently, researchers developed a transcranial brain atlas (TBA) based on the balloon inflation model, which directly maps brain atlas labels onto the scalp space (Xiao et al., 2018). This TBA can be used to both optimize optode montages and determine the anatomical origins of fNIRS signals (Xiao et al., 2018; Zhao et al., 2021). The performance of TBA-guided optode arrangement has been validated, showing significant improvements compared to the traditional 10-20 system-assisted optode arrangement (Jiang et al., 2020; Xiao et al., 2018). While the balloon inflation model-based TBA has made progress in fNIRS spatial localization and offers a straightforward approach, it does not fully account for the complex nature of light propagation in the head during fNIRS imaging. Specifically, the balloon inflation model oversimplifies the measured volume by representing it as a single cortical point, which fails to comprehensively capture the spatially distributed cortical regions that fNIRS can detect (Cai et al., 2021).

The photon measurement density function (PMDF) has emerged as a novel approach to more accurately describe the spatial distribution detected by fNIRS channels (Brigadoi et al., 2018; Vera et al., 2022). PMDF not only identifies the banana-shaped spatial distribution that contributes to the channel measurement signal but also precisely provides the sensitivity of the measurement signal to absorption changes at each location within this spatial distribution (Arridge and Schweiger, 1995). With PMDF, the accuracy and reliability of fNIRS spatial localization can be significantly improved. Strictly, calculating PMDF requires individual magnetic resonance imaging (MRI) images of subjects to construct realistic head models and perform Monte Carlo simulations (Machado et al., 2018). However, MRI scans are not routinely performed in fNIRS studies, as they clearly conflict with the economic advantages and convenience of fNIRS (Tsuzuki and Dan, 2014).

To address the lack of individual MRI scans, researchers have proposed using standard MRI, such as Colin27 and MNI152 in the MNI space, to construct head models and calculate PMDF. These methods have been employed to guide the design of optode arrangement s in fNIRS experiments (Brigadoi et al., 2018; Wijeakumar et al., 2015; Zimeo Morais et al., 2018) and localize anatomical regions of fNIRS data (Aasted et al., 2015; Cai et al., 2021). Although this approach circumvents the need for individual MRI scans, it has two significant limitations. First, there are inevitable differences between the standard head models and the actual head anatomical structures of individual subjects (Fu and Richards, 2021), with the extent of these differences varying from person to person. Currently, there is a lack of quantitative, large-sample studies assessing the impact of these anatomical differences on localization accuracy. Second, although group-averaged PMDF using large-sample MRI has gained attention (Fu and Richards, 2021), the predefined channel space is limited to channels whose probes are located in the 10-10 system and meet specific source-detector distances (e.g., 1-5 cm). While this channel space is convenient for generalization across different subjects, it is highly restrictive (Brigadoi et al., 2018).

To address the above challenges, this study proposed a continuous fNIRS standard channel space and a PMDF-based transcranial brain atlas (PMDF-TBA) derived from a large-sample MRI dataset. We began by defining a triple-parameter standard channel space that allowed for flexible description of any fNIRS channel, based on its channel center positions (of source and detector), source-detector orientations, and source-detector distance. Using this foundation, we conducted Monte Carlo photon propagation simulations on MRI images of 48 healthy adults. These simulations calculated the group-averaged sensitivity of channels to gray matter and brain regions of three atlases, formed the PMDF-TBA. Furthermore, we use leave-one-out cross-validation (LOOCV) to quantitatively evaluate the predictive ability of PMDF-TBA for the sensitivity of unseen individuals. We compare its performance with that of traditional PMDFs calculated based on single Colin27 or MNI152 head models.

## 2. Theory and methods

### 2.1 Standard channel space

We propose a standard channel space for fNIRS that employs three parameters: **s**, θ, and d, to describe any channel on the head surface. The first parameter, **s**, represents the midpoint position of the curve connecting the emitter and detector optodes on the scalp (Fig. 1A). It is based on the proportional coordinates (*p*_*NZ*_, *p*_*AL*_) of the previously proposed continuous proportional coordinate (CPC) space (Xiao et al., 2018). The coordinate 0 < *p*_*NZ*_ < 1 indicates the channel center’s position in the anterior-posterior direction, with smaller *p*_*NZ*_ values representing more anterior locations. The coordinate 0 < *p*_*AL*_ < 1 indicates the channel center’s position in the left-right direction, with smaller *p*_*AL*_ values representing more leftward locations.

**Figure 1.**
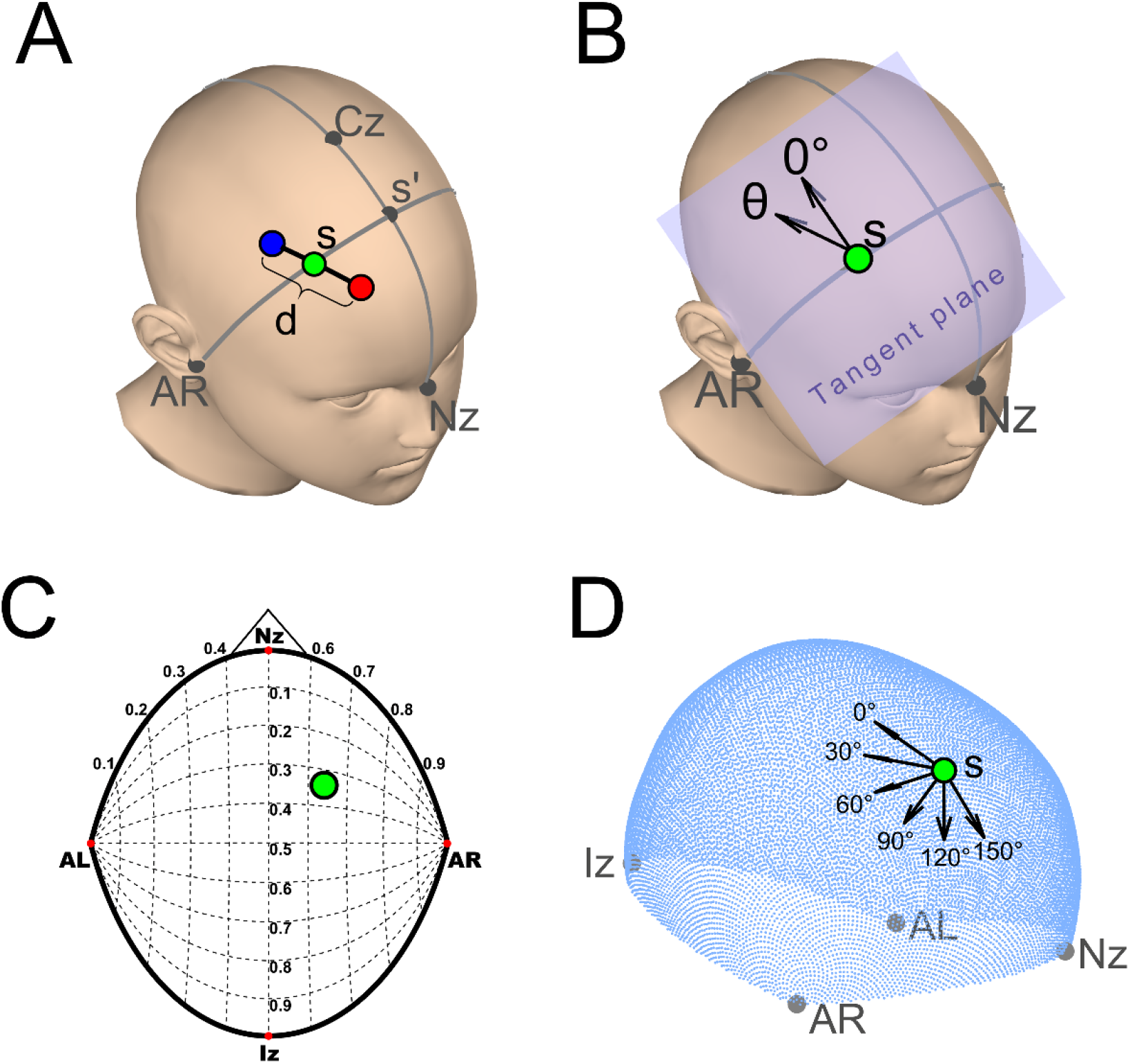
The channel space and an example channel (**s**=[0.34,0.64], **θ**=30°, **d**=3cm). (A) Definition of parameters **s** and **d**: The center **s** (green dot) of the channel is determined by a reference line (Nz-Cz-Iz) and an active line (AL-s-AR). The reference line is the intersection of the scalp surface with the sagittal plane through Nz, Cz, and Iz, while the active line is the intersection of the scalp surface with the plane passing through AL, AR, and **s**. The reference and active lines intersect at point s’. The proportional coordinates *p*_*NZ*_ and *p*_*AL*_ are calculated as *p*_*NZ*_ = *L*_*NZ*−s′_/*L*_*NZ*−s′−IZ_ and *p*_*AL*_ = *L*_*AL*−s_ /*L*_*AL*−s−AR_. The parameter **d** represents the distance between source (red dot) and detector (blue dot) along the head surface (black line). (B) Definition of parameter **θ**: **θ** is the angle between the channel orientation vector and the 0° direction vector, both of which lie on the tangent plane of point **s**. The 0° direction is defined by a unit vector perpendicular to the active line (AL-s-AR), pointing backward from **s**. (C) The channel center **s** is projected onto a 2D plane for visualization purposes. (D) 18,000 points (CPC2mm) are sampled from the continuous proportional coordinate (CPC) space to serve as the channel center points **s**, and 6 channel orientations are sampled for each center **s**.

The second parameter, **θ**, describes the channel’s orientation (Fig. 1B), which is the angle between the direction from the source to the detector and the predefined zero-direction vector on the tangent plane of **s**. This orientation parameter is derived from the previously proposed scalp-measurement based parameter (SGP) space (Jiang et al., 2022). The third parameter, **d**, is the length of the curve connecting the source and detector optodes on the scalp (Fig. 1A). These parameters establish a one-to-one mapping to any channel on a subject’s head. Using the BNU projection method (Xiao et al., 2018), the channel center parameter s can be transformed onto a two-dimensional plane for visualization (Fig. 1C).

To facilitate computation and practical application, we discretized the channel space. Specifically, we uniformly sampled 18,000 CPC points in the CPC space as a point set of channel center **s** (Fig. 1D), with a distance of approximately 2 mm between adjacent points. This point set was referred to as CPC2mm. Considering that the PMDF remains unchanged when the positions of the source and detector were exchanged, the range of the channel orientation **θ** was set to 0°-180°, with uniform sampling at 30° intervals, i.e., {θ_1_=0°, θ_2_=30°, θ_3_=60°, θ_4_=90°, θ_5_=120°, θ_6_=150°}. The distance **d** can also be discretized into several different lengths, such as {20 mm, 25 mm, 30 mm, 35 mm, 40 mm}. As this study used adult MRI data, we used **d** = 30 mm as an example to introduce the construction of the PMDF-TBA. This discretization process resulted in a total of 108,000 channels evenly distributed across the scalp surface.

### 2.2 Individual-level channel-wise sensitivity for gray matter and brain regions of atlas

The individual-level sensitivity of a channel for gray matter and brain regions of atlas depends on the PMDF. To obtain the PMDF of the channel, we first constructed subject-specific head models using structural magnetic resonance imaging (sMRI) data from 48 adult subjects, randomly selected from the Southwest University Adult Lifespan Dataset (Wei et al., 2018). The demographic data and scanning parameters were provided in the supplementary materials. The head models were constructed using the headreco command in the SimNIBS toolbox (Saturnino et al., 2019). Each head model included five tissue types: scalp, skull, cerebrospinal fluid, gray matter, and white matter.

Given a set of channel (s_*i*_, θ_*j*_, *d*=3*cm*), hereafter referred to as (s_*i*_, θ_*j*_), and a subject head model, we can identify the positions of the source and detector optodes. This was achieved by rotating the 0°direction vector of s_*i*_ around its normal vector by θ_*j*_ degrees, resulting in a channel orientation vector. This channel orientation vector and the normal vector of s_*i*_ formed a plane. The intersection of this plane with the scalp surface created an intersection curve. All the intersection points on this curve belonged to the CPC2mm point set. Finally, we searched along both directions of the intersection curve to find two intersection points with geodesic distances closest to 15 mm from s_*i*_, which served as the source and detector.

The PMDF of the channel (s_*i*_, θ_*j*_) was calculated using the Adjoint Monte Carlo method (Custo et al., 2010):

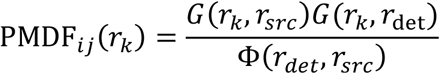

where *r*_*k*_ represented the location (vertex) in the individual space, *G*(*r*_*k*_, *r*_*src*_) and *G*(*r*_*k*_, *r*_*det*_) were the Green’s functions at *r*_*k*_, which were generated by light propagation from the source *r*_*src*_ and detector *r*_det_, respectively. Φ(*r*_*det*_, *r*_*src*_) denoted the photon fluence at *r*_*det*_ , originating from *r*_*src*_. PMDF_*ij*_(*r*_*k*_) represented the sensitivity of the channel to the unit volume at location *r*_*k*_. The PMDF can be used to calculate the maximum sensitivity position of the channel within the brain:

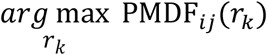

The above simulation was performed in the Mesh-based Monte Carlo toolbox (Fang, 2010), and optical properties (Supplementary Table 1) of the five tissues in the head model were the average optical properties of four commonly used near-infrared wavelengths (690, 750, 780, and 830 nm) (Strangman et al., 2013).

To calculate the sensitivity values for gray matter and brain regions, we first converted the PMDF values from vertex-based to tetrahedral-based representation:

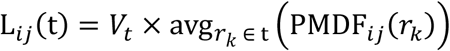

where **t** represented the t-th tetrahedron, and *V*_*t*_ represented the volume of that tetrahedron. We then calculated the sensitivity of channel (s_*i*_, θ_*j*_)to the gray matter:

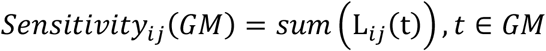

Given the **m**-th brain region in an atlas as the ROI, we transformed it into the individual space and calculated the sensitivity and specificity of the channel to that ROI.

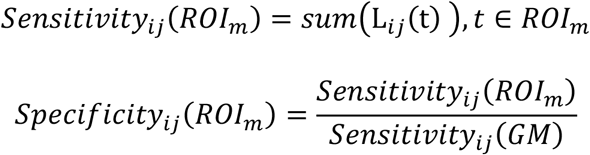

We used brain regions from three widely adopted anatomical atlases: Brodmann areas (BA), the 2nd edition Automated Anatomical Labeling Atlas (AAL2), and the LONI (Laboratory of Neuro Imaging, University of Southern California) Probabilistic Brain Atlas constructed from 40 adult humans (LPBA40).

### 2.3 Group-level channel-wise sensitivity for gray matter and brain regions of atlas

At the group level, the average sensitivity and variability of channel (s_*i*_, θ_*j*_) to gray matter can be obtained through aggregate analysis of individual data:

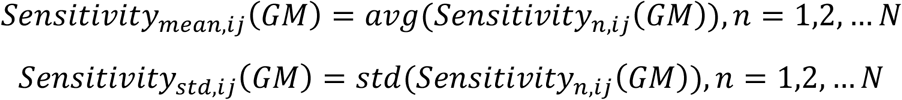

where **n** represented the **n**-th subject and **N** was the number of subjects. By iterating through all channels, we obtained the gray matter sensitivity map of PMDF-TBA. Similarly, we calculated the average sensitivity and variability of channel (s_*i*_, θ_*j*_) to the **m**-th ROI of the brain atlas at the group level:

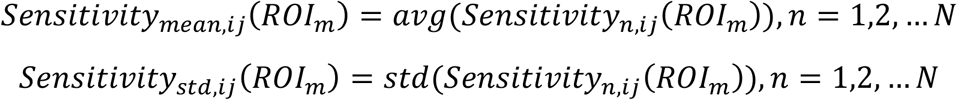

By iterating through all channels, we obtained the sensitivity map (mean or standard deviation) for the *ROI*_*m*_ . Additionally, we calculated the average specificity and its variability of channel (s_*i*_, θ_*j*_) to brain atlas regions at the group level:

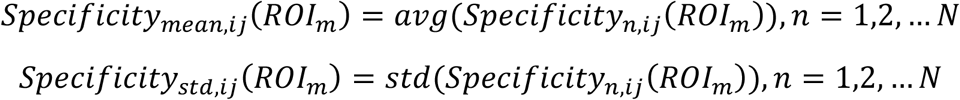

By iterating through all channels, we obtained the specificity map (mean or standard deviation) of PMDF-TBA for the *ROI*_*m*_.

Considering that the PMDF distribution of channel (s_*i*_, θ_*j*_) may cover multiple Brodmann areas, we further calculated the brain region label with maximum sensitivity:

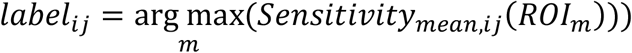

thereby obtaining the maximum sensitivity label map of PMDF-TBA. The sensitivity and specificity values corresponding to the maximum sensitivity label formed the maximum sensitivity map and maximum specificity map, respectively. For ease of observation, we averaged the sensitivities of channels in 6 different orientations and obtained the label with maximum average sensitivity.

To obtain the brain location with maximum sensitive for channel (s_*i*_, θ_*j*_) at the group level, we transformed the coordinates of individual maximum positions from individual space into MNI space and then calculated the group average coordinates. The degree of variability across subjects can be obtained by calculating the average distance from individual positions to the group mean position in standard space. Although the maximum sensitivity position and its variability were computed as part of the PMDF-based metrics, as they were not the primary focus of this study, they would not be explicitly reported in the Results section.

### 2.4 Validation of PMDF-TBA

To assess the predictive ability of PMDF-TBA on unseen subjects, we employed a LOOCV approach. Specifically, 47 subjects were selected from the total of 48 subjects to construct PMDF-TBA47, while the remaining subject served as the test subject. The ground truth for the test subject was the sensitivity of channels to the gray matter or brain regions of BA, as well as the maximum sensitivity labels, which were derived from his or her structural image. The error metrics for sensitivity were absolute error and correlation coefficient; the error metric for maximum sensitivity brain atlas labels was the proportion of incorrectly predicted channels relative to the total number of channels.

For comparison, we also constructed PMDF-TBA using two standard head MRI (Colin27 and MNI152) commonly used by previous researchers (Fu and Richards, 2021), and obtained PMDF-TBA-Colin27 and PMDF-TBA-MNI152, respectively. Similarly, we employed LOOCV to predict the ground truth of the test subject. Finally, paired-sample t-tests were used to compare the predictive performance of the PMDF-TBA47 with PMDF-TBA-Colin27 and PMDF-TBA-MNI152.

Additionally, we defined an effective channel range, 0.15 ≤ *p*_*nz*_ 0.85,0.15 ≤ *p*_*al*_ ≤ 0.85, to avoid optodes of channels being placed outside the boundaries of the CPC system or on the ears and eyes. In all validations, we only evaluated channels whose centers were within this range. Furthermore for sensitivity map, we noticed that some channels within effective range still had small sensitivity values, which may underestimating the prediction error of sensitivity values. Therefore, we only considered channels with sensitivity values greater than the 5th percentile of the maximum value in the sensitivity map (either the ground truth or PMDF-TBA47).

## 3. Results

### 3.1 Constructed PMDF-TBA

The constructed PMDF-TBA is shown in Figure 2, including sensitivity maps (Fig. 2A-D) and maximum sensitivity label map for the Brodmann areas (Fig. 2E). Figure 2A displays the sensitivity map of channels to gray matter, with channels covering the bilateral frontal lobes exhibiting the highest sensitivity, while channels covering the anterior temporal lobe and longitudinal fissure of the parietal lobe show lower sensitivity. This sensitivity distribution is somewhat related to the distribution of the brain-to-scalp distance (Supplementary Figure 1): channels with higher sensitivity tend to have shorter brain-to-scalp distances, while channels with lower sensitivity have longer distances. Figure 2B presents the sensitivity map for the premotor cortex (BA 6), with channels in the central region having the highest sensitivity and those in the peripheral regions showing relatively lower sensitivity. Figure 2C illustrates the sensitivity maps for the primary motor cortex (BA 4), revealing that channel orientation influences sensitivity. Channels with a same center position exhibit higher sensitivity when oriented parallel to the long axis of BA 4 compared to those oriented perpendicular to the long axis. Figure 2D illustrates that for the pars opercularis of Broca’s area (BA 44), the channels with maximum sensitivity across different orientation maps have similar channel center positions. Additionally, the supplementary material provides the variation of sensitivity across subjects (Supplementary Figure 2) and the group-averaged specificity (Supplementary Figure 3).

**Figure 2.**
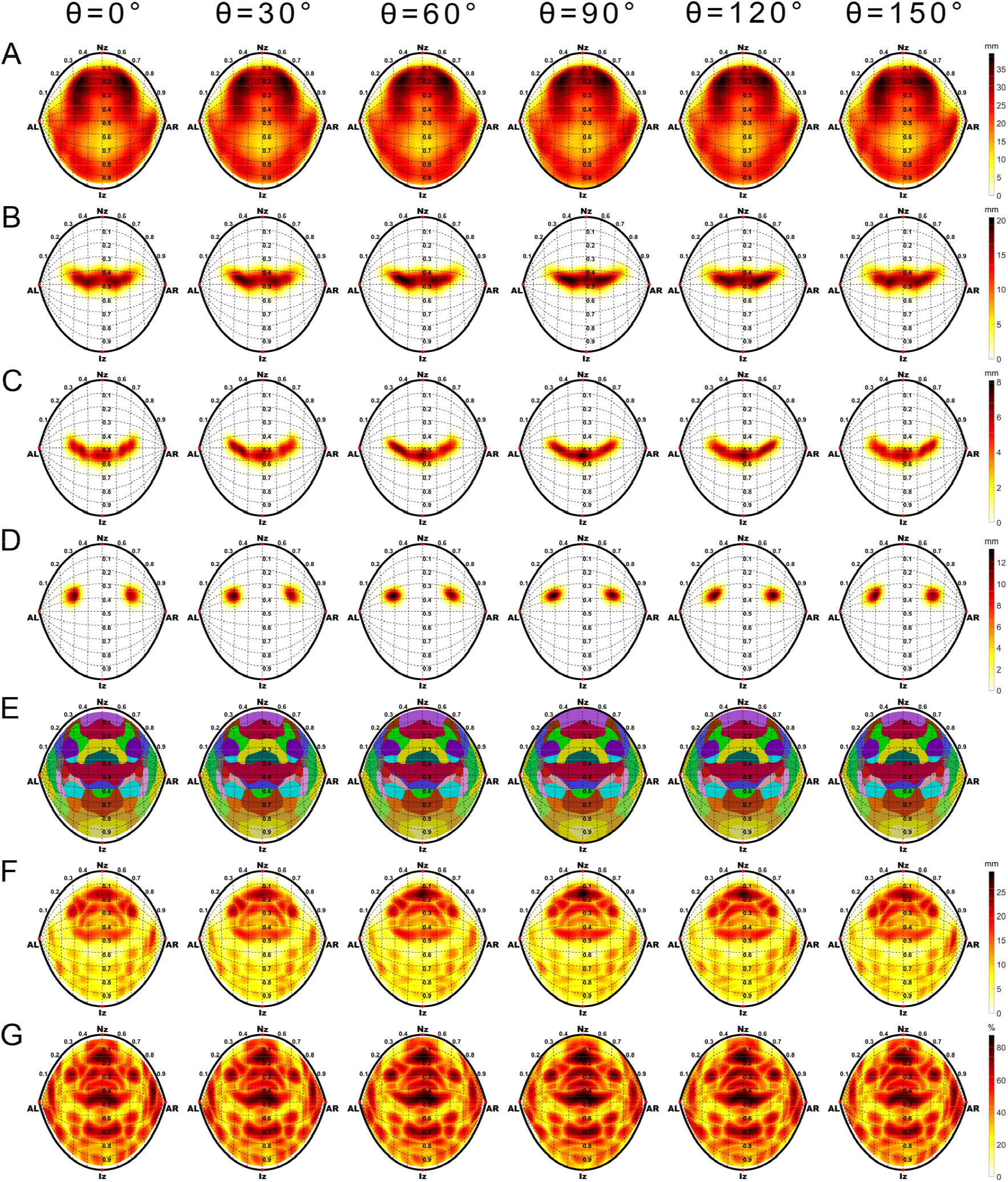
The PMDF-TBA. (A) Sensitivity map of gray matter. (B-D) Sensitivity maps of Brodmann areas: BA 6 (B), BA 4 (C), and BA 44 (D). (E) Maximum sensitivity label map of the Brodmann areas; the corresponding label names for each color are provided in Supplementary Figure 8. (F) Maximum sensitivity map. (G) Maximum specificity map.

Figure 2E displays the maximum sensitivity label map based on the Brodmann areas. Interestingly, we found that even when multiple channels had same maximum sensitivity label, their maximum sensitivity (Fig. 2F) or specificity (Fig. 2G) may differ. Moreover, in the maximum sensitivity map (Fig. 2F), brain regions in the frontal lobe exhibit the highest sensitivity, corresponding to the shorter brain-to-scalp depth in this area. PMDF-TBA for AAL2 and LPBA40 are provided (Supplementary Figures 4-5) for further reference.

Figure 3 presents the orientation-averaged PMDF-TBA for three widely adopted anatomical atlases. For the Brodmann areas (Fig. 3A), we provide the maximum orientation-averaged sensitivity label map (Fig. 3B). Each point in this map is obtained by averaging the sensitivity values of 6 channels with different orientations but the same channel center position, and then assigning the brain atlas label with the highest orientation-averaged sensitivity. Figures 3C and 3D show the corresponding maximum orientation-averaged sensitivity and maximum orientation-averaged specificity, respectively. Similarly, we also present the corresponding results for the AAL2 (Fig. 3E-H) and the LPBA40 (Fig. 3I-L).

**Figure 3.**
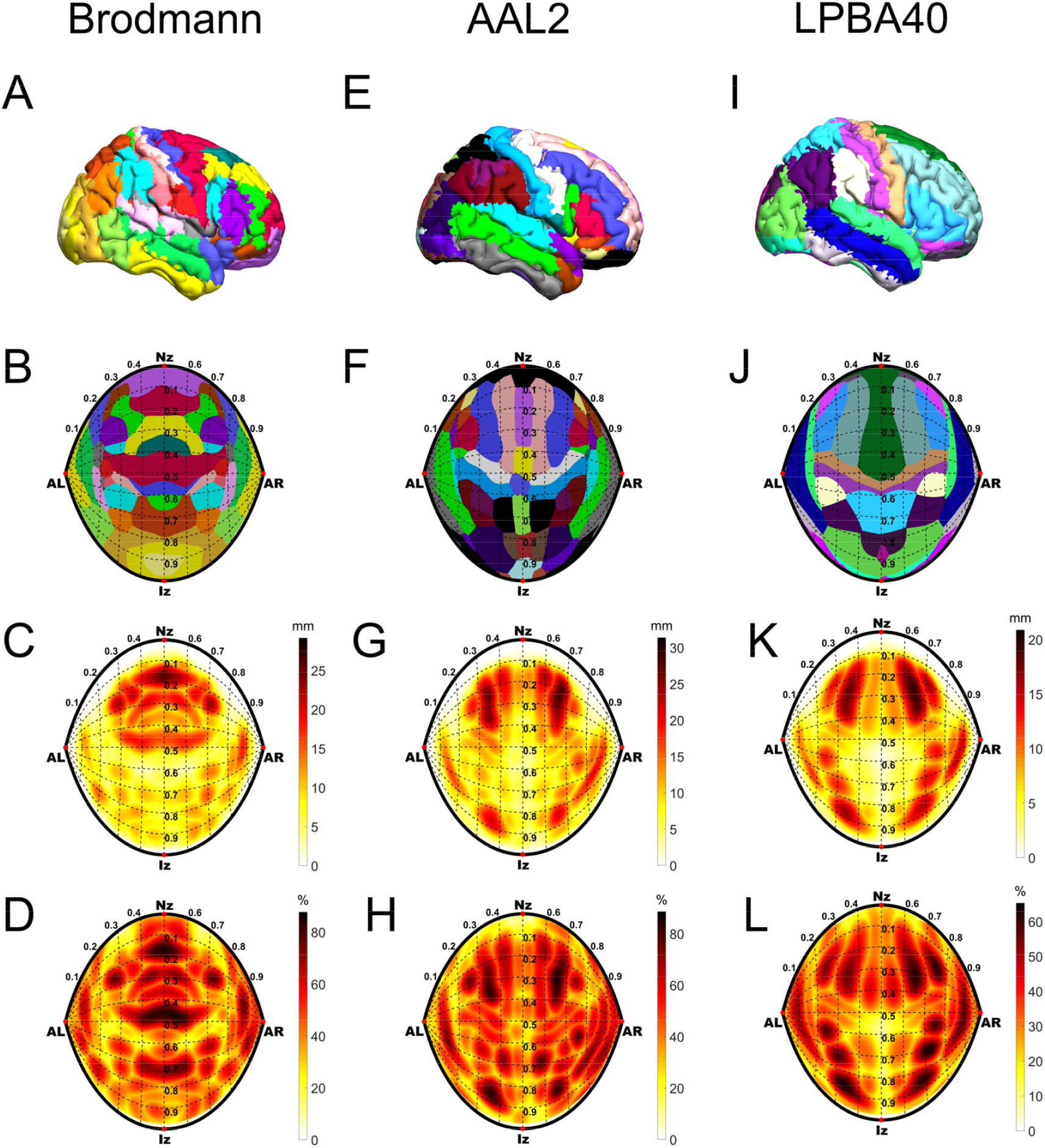
Maximum orientation-averaged sensitivity labels, maximum sensitivity, and maximum specificity for three atlases. (First row) The three atlases: Brodmann areas (first column), AAL2 (second column), and LPBA40 (third column). (Second row) Maximum orientation-averaged sensitivity labels for each atlas. (Third row) Corresponding maximum sensitivity for each atlas. (Fourth row) Corresponding maximum specificity for each atlas.

### 3.2 Results of Prediction

For the prediction of gray matter sensitivity, Figure 4A shows the ground truth of a randomly selected subject, PMDF-TBA47 from the remaining 47 subjects and the mean absolute error (MAE) of 48 times of LOOCV. Across all subjects, the MAE was 4.45 ± 1.4 mm, and the average correlation coefficient reached 0.81 ± 0.07. Regarding Brodmann area sensitivity, except for 7 regions that could not be effectively evaluated due to their deep locations and sensitivities below the computational precision, the remaining 41 regions had a MAE of 0.85 ± 0.20 mm and an average correlation coefficient of 0.84 ± 0.05 for PMDF-TBA predictions. Taking BA6, BA4, and BA44 as examples (Fig. 4B-D), their average MAEs were 1.99 ± 0.74 mm, 0.88 ± 0.33 mm, and 1.24 ± 0.36 mm, respectively, while their average correlation coefficients reached 0.91 ± 0.06, 0.87 ± 0.09, and 0.90 ± 0.06, respectively.

**Figure 4.**
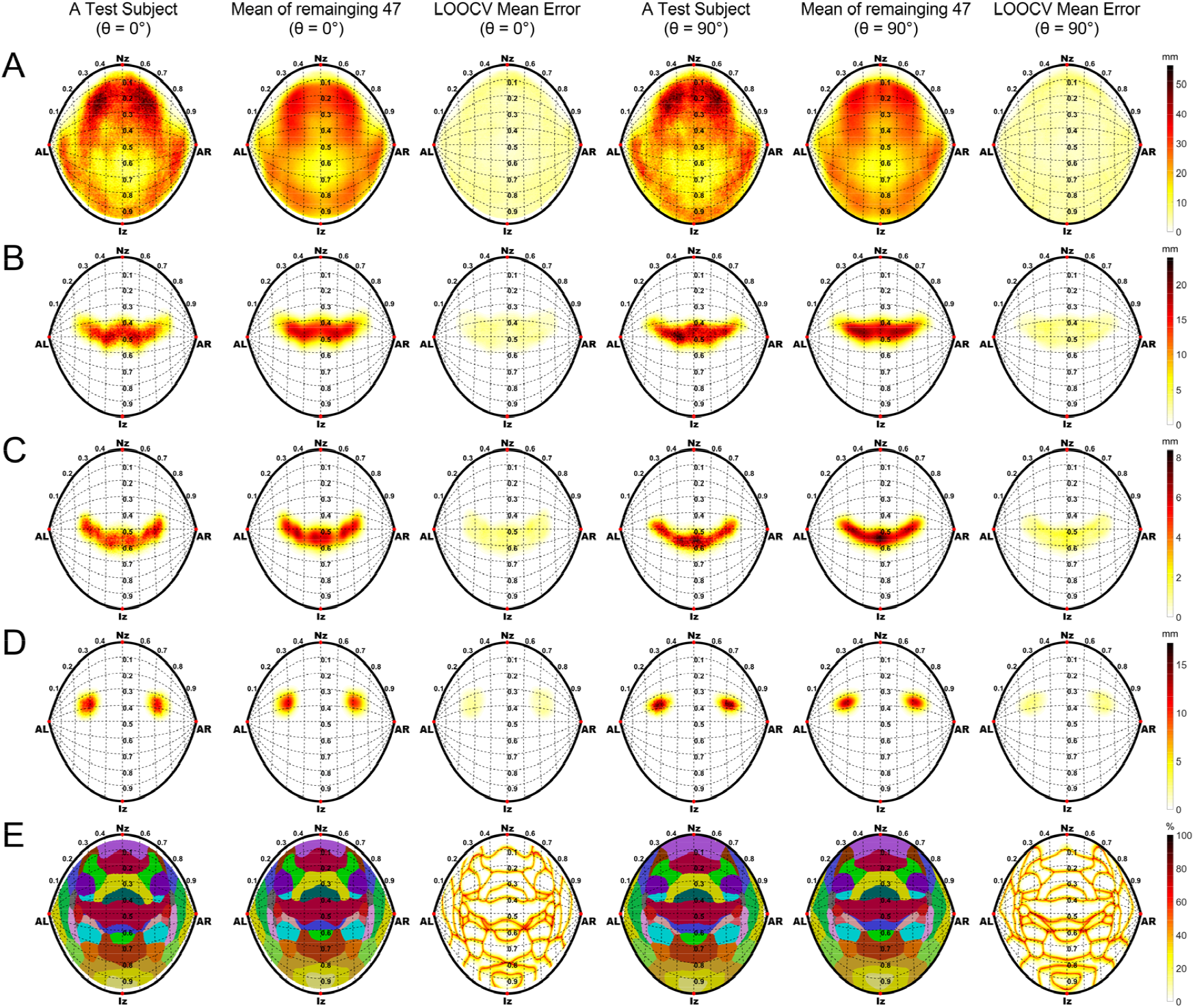
Prediction performance evaluation using leave one out cross-validation. Sensitivity maps of gray matter (A), BA 6 (B), BA 4 (C) and BA 44 (D); Maximum sensitivity label maps of Brodmann areas(E). Ground truth map (first and fourth column), predicted map (second and fifth column) and prediction error (third and sixth column).

In terms of predicting maximum sensitivity labels, Figure 4E presents the ground truth for a randomly selected subject, PMDF-TBA47, and the average prediction error of LOOCV. The average prediction error rate of PMDF-TBA across all channels was only 16.5%. We found that within the TBA regions of interest (ROIs), the prediction error rates were low, with higher error rates only occurring at ROI boundaries.

Figure 5 shows the prediction errors of sensitivity and maximum sensitivity labels for unseen subjects using PMDF-TBA constructed based on two standard head models (Colin27 and MNI152). For sensitivity prediction, the MAEs of PMDF-TBA-Colin27 and PMDF-TBA-MNI152 were 1.51 ± 0.24 mm and 1.75 ± 0.28 mm, respectively, both significantly larger than PMDF-TBA47 (Colin27: paired t-test, t=17.32, df=47, p<0.0001; MNI152: paired t-test, t=18.50, df=47, p<0.0001). Moreover, the average correlation coefficients of the PMDF-TBA-Colin27 and PMDF-TBA-MNI152 were 0.68 ± 0.05 and 0.64 ± 0.05, respectively, also significantly lower than PMDF-TBA47 (Colin27: paired t-test, t=-26.10, df=47, p<0.0001; MNI152: paired t-test, t=-49.54, df=47, p<0.0001).

**Figure 5.**
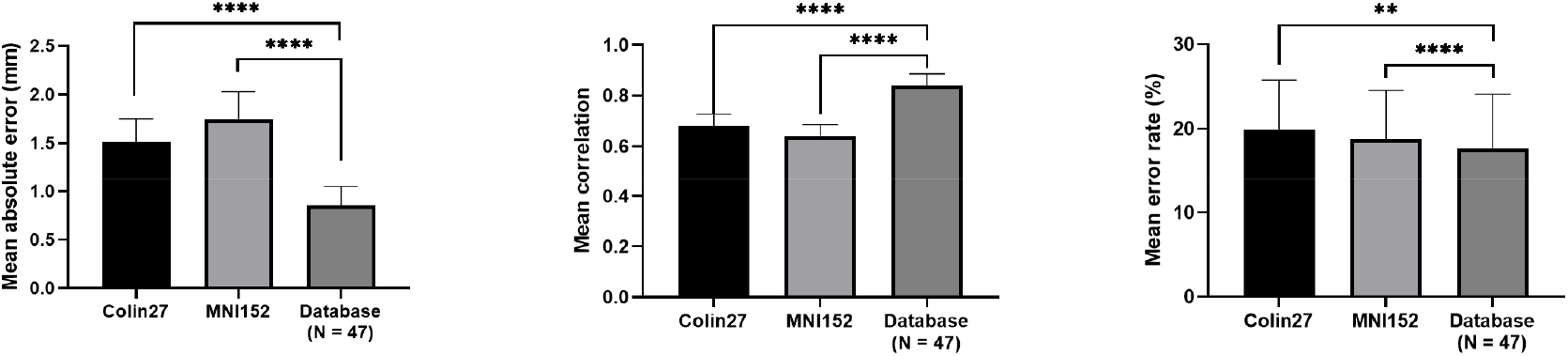
Prediction errors of unseen subjects by PMDF-TBA constructed using Colin27 (PMDF-TBA-Colin27), MNI152 (PMDF-TBA-MNI152), and a large sample MRI database with 47 subjects (PMDF-TBA47). The error metrics were (A) mean absolute error, (B) mean correlation coefficient, and (C) mean error rate. **** indicates that the prediction errors of the two TBAs in the same test subjects (N=48) reached p<0.0001 through a two-tailed paired t-test, while ** denotes p<0.01 for a two-tailed test.

Regarding the prediction of maximum sensitivity labels, the average error rates of PMDF-TBA-Colin27 and PMDF-TBA-MNI152 were 19.9 ± 5.8% and 18.8 ± 5.8%, respectively, which were also significantly higher than PMDF-TBA47 (Colin27: paired t-test, t=3.45, df=47, p=0.0011; MNI152: paired t-test, t=4.76, df=47, p<0.0001). These results demonstrate that PMDF-TBA47 exhibits significantly better predictive performance compared to the two standard head models.

## 4. Discussion

In this study, we proposed a standard channel space for fNIRS defined by triple parameters (**s, θ, d**). Within this space, we collected discrete channels and performed photon propagation simulations using a large MRI dataset. We obtained sensitivity metrics for gray matter and brain regions of three atlases, as well as the maximum sensitivity labels, forming the PMDF-TBA. By comparing the performance of PMDF-TBA constructed using the large dataset and single standard head models in predicting sensitivity and maximum sensitivity labels of unseen individuals, we found that the PMDF-TBA constructed using the large dataset exhibited superior predictive ability. Here we discuss the advantages of the standard channel space, the effectiveness and stability of the group PMDF-TBA, the potential applications of PMDF-TBA, and the similarity of PMDF-TBA with different orientations. Finally, we analyze the limitations of this study and provide an outlook on future work.

### 4.1. Advantages of the Standard Channel Space

The standard channel space offers several advantages. Firstly, it has continuity, allowing the position of any channel to be described. This facilitates the intuitive display of the distribution of channel characteristics, such as sensitivity distribution. It also enables the description of arbitrary optode arrangements, which providing a broad optimization space for optode arrangement design, and accurate description of optode arrangements, which offers convenience for subsequent researchers to replicate experiments. Secondly, channels across different subjects described by the same set of parameters have correspondence. By mapping each subject’s channels to this unified space, we can conveniently aggregate and compare information of the corresponding channel at the group level. Lastly, researchers can manually measure the position of channels on the scalp based on the three parameters without relying on specialized positioning devices, offering practicality and convenience.

The standard channel space explicitly represents key parameters of channels, including center position, orientation, and source-detector distance. In contrast, the literature often describes the channel location by defining two optodes on 10-10 system (or its derivatives) points and forms channels which meet certain distance ranges (e.g., 1-5 cm) (Brigadoi et al., 2018; Fu and Richards, 2021; Zimeo Morais et al., 2018). Although this traditional method is simple and straightforward, it does not fully utilize the key geometric parameters of channel positions, leading to several issues.

Firstly, the lack of a channel center position parameter makes it impossible to accurately present the distribution of characteristic values. For example, researchers can only represent the averaged sensitivity values of multiple adjacent channels within a local region at a single optode position (Strangman et al., 2014; Whiteman et al., 2018). This approach ignores the differences in position, orientation, and source-detector distance of different channels, greatly reducing the accuracy of sensitivity values. Secondly, there is limited research on the influence of orientation factors on sensitivity (Cai et al., 2021), possibly due to the lack of channel orientation parameters. However, the orientation factor cannot be ignored because the banana-shaped photon migration path formed by channels has anisotropy, resulting in different sensitivities of channels with different orientations to the underlying cortex. Thirdly, the lack of a source-detector distance parameter is not conducive to establishing consistent channels across subjects. For example, channels formed by a pair of 10-10 points as source and detector optodes have different distances in subjects with different head circumference sizes, while many devices adopt fixed distance settings (e.g., 3 cm for adults). The standard channel space addresses these issues by introducing key geometric parameters, providing a more refined and flexible analysis framework for fNIRS research.

### 4.2. Effectiveness of PMDF-TBA

The effectiveness of PMDF-TBA lies in its comprehensive characterization of the fNIRS imaging process to obtain localization information. Compared to the localization information reported by balloon inflation model-based TBA, which includes the MNI coordinates of the cortical surface corresponding to the channel center and the probability of these cortical surface points falling into different brain atlas parcellations (Xiao et al., 2018), PMDF-TBA not only reports the MNI coordinates of the maximum sensitivity on the cortical surface corresponding to each channel but also provides the detection sensitivity and specificity for different brain atlas parcellations. These differences reflect the fundamental distinctions between the two methods in describing the fNIRS imaging process: the balloon inflation model only approximates the most sensitive point on the cortical surface detected by the channel, while PMDF can fully characterize the sensitivity distribution over the entire imaging range, which better aligns with the physical imaging process of fNIRS. Therefore, although balloon inflation model-based TBA has strong generalizability and is applicable to various transcranial techniques such as fNIRS and transcranial magnetic stimulation, PMDF-TBA can provide more comprehensive and refined imaging information specific to fNIRS technology, thus having unique advantages in solving localization problems.

Additionally, the effectiveness of PMDF-TBA is also demonstrated by its predictive ability for unseen subjects. Due to the lack of individual structural images, PMDF is usually simulated using standard head models (such as Colin27 and MNI152) (Aasted et al., 2015; Brigadoi et al., 2018; Zimeo Morais et al., 2018). However, at the theoretical level, PMDFs derived from single standard head models may deviate from the group average, thus having limited representativeness. In contrast, PMDF calculated based on a group should be more representative. To verify this theoretical inference, we experimentally evaluated and compared the generalization ability of PMDF-TBA constructed based on single standard head models and PMDF-TBA based on a large dataset. The results showed that the generalization ability of PMDF-TBA based on a large dataset was significantly better than that of PMDF-TBA based on single standard head models. This experimental finding is consistent with the theoretical inference, further supporting the conclusion that our PMDF-TBA based on a large dataset has high group representativeness.

### 4.3. Stability of PMDF-TBA

When constructing PMDF-TBA based on MRI database, the size of the database is crucial to its generalization ability. If the database is too small, PMDF-TBA may overly rely on individual characteristics, lacking sufficient group representativeness and leading to insufficient generalization ability. However, when we increase the database size to a point where the generalization ability of PMDF-TBA reaches a stable state, further increasing the data volume often brings very limited improvement in generalization ability but consumes a large amount of storage and computational resources. Therefore, we need to assess whether the predictive ability of the current PMDF-TBA has reached a stable state to achieve a balance between generalization ability and resource efficiency.

To evaluate the stability of PMDF-TBA, we constructed multiple PMDF-TBAs using varying database sizes (from 1 to 25 with a step size of 1) and evaluated their prediction accuracy of the sensitivity of brain regions for three commonly used atlases (Brodmann, AAL2, and LPBA40) on unseen subjects (details provided in the supplementary materials). The results (supplementary Figure 6) showed that as the database size increased, the generalization ability of PMDF-TBA first increased rapidly and then stabilized when the number of subjects exceeded 6. This indicates that our PMDF-TBA model constructed based on 48 subjects has reached a stable state and has good generalization ability.

### 4.4. Potential applications of PMDF-TBA

PMDF-TBA possesses knowledge attributes, providing valuable information about the sensitivity of arbitrary channels to ROIs, which helps researchers gain an understanding of the distribution characteristics of sensitivity. Based on this knowledge attribute, PMDF-TBA can be applied to solve spatial localization problems. Before data acquisition, researchers can utilize 3D digitization systems in conjunction with the orientation-averaged sensitivity map provided by PMDF-TBA (supplementary Figure 7) to identify channel positions that are highly sensitive to ROIs, thereby optimizing optode arrangements. After data acquisition, by recording the position of the optode arrangements on the subject’s scalp using a 3D digitization system, PMDF-TBA can provide comprehensive localization information for the optode arrangement, including the MNI coordinates of the maximum sensitivity for each channel and the sensitivity and specificity of the channels to each brain atlas parcellation. This process does not require the subject’s structural images or complex photon propagation simulations.

Furthermore, PMDF-TBA can be incorporated in post-experiment data analyses. When converting fNIRS light intensity signals into hemoglobin signals, querying the corresponding sensitivity in PMDF-TBA as the optical path length can improve the accuracy of the conversion (Strangman et al., 2003). The other application is in ROI analysis, researchers often need to aggregate multiple channels for a single ROI (Sun et al., 2018; Yanagisawa et al., 2010). With the aid of prior information provided by PMDF-TBA, researchers can perform a weighted average of multiple channels’ response values to the ROI with sensitivity as the weight, obtaining a comprehensive response index and improving the accuracy of the analysis.

### 4.5. Similarity of PMDF-TBA with different orientations

Through quantitative analysis, we found that channels located at the same center position but with different orientations exhibit a certain degree of similarity in their sensitivity to brain regions. This similarity is reflected in the high correlation between the sensitivity maps of PMDF-TBA for the same Brodmann areas across six different orientations, with an average correlation coefficient as high as 0.96±0.05. Given this similarity, to simplify the analysis and facilitate observation, we averaged the sensitivity maps of different orientations. The resulting averaged PMDF-TBA integrates information from multiple orientations, allowing researchers to optimize optode arrangements and understand the spatial distribution of sensitivity on this basis.

### 4.6. Limitations & Further work

This study has some limitations. First, PMDF-TBA currently only adopts the commonly used 3cm as the source-detector distance. In the future, PMDF-TBA for arbitrary channel distances can be constructed based on the construction framework described in this paper. Second, the current PMDF-TBA can only report the sensitivity of channels to gray matter and regions from three widely adopted anatomical brain atlases, but this method can be extended to any anatomical (Desikan et al., 2006; Fan et al., 2016; Schaefer et al., 2017) and functional brain atlases (Jiang et al., 2020). Third, the current PMDF-TBA is constructed based on MRI of healthy adults, and it is unclear whether it can be applied to other populations. In the future, PMDF-TBA suitable for different age groups (Fu and Richards, 2021; Zhang et al., 2021) and populations with mental disorders can be constructed. Fourth, the current PMDF-TBA calculates group-averaged sensitivity for group experiments, but for individual studies, individual PMDF-TBA can be constructed based on individual MRI to achieve individualized localization(Machado et al., 2018, 2014). Fifth, the current application of PMDF-TBA to optode arrangement design relies on visual assistance. In the future, computer-aided automatic optimization of optode arrangement design can be adopted to fully utilize the sensitivity information provided by PMDF-TBA and customize the optimal optode arrangement (Zhao et al., 2021).

In PMDF-TBA, PMDF is the key to calculate the sensitivity of arbitrary ROIs. In addition, PMDF can also be used for image reconstruction in diffuse optical tomography (Custo et al., 2010). However, this study did not store the PMDF of each channel because the amount of PMDF data for each channel is large (about 35 Mb), and storing 108,000 channels for all subjects (N=48) would require enormous storage resources (about 173 TB). Therefore, how to calculate and store the group PMDF of arbitrary channels is the next problem to be solved.

## Author contributions section

Lijiang Wei: Methodology, Investigation, Visualization, Writing. Yang Zhao: Methodology, Investigation, Writing. Farui Liu: Methodology. Yuanyuan Chen: Writing. Yilong Xu: Methodology. Zheng Li, Writing. Chaozhe Zhu: Conceptualization, Methodology, Supervision, Writing

## Acknowledgements

This work was sponsored by the National Natural Science Foundation of China (82071999 and 61431002). The authors declare no competing financial interests.

