## Supplementary for "Transcranial Brain Atlas Based on Photon Measurement Density Function in a Triple-Parameter Standard Channel Space"

### 1. Participants and Image Acquisitions

Structural images of 48 subjects were randomly selected from the cross-sectional Southwest University adult lifespan dataset. The subjects had a mean age of 20.0±1.2 years, and 24 of them were female. High-resolution T1-weighted anatomical images were obtained using a magnetization-prepared rapid gradient echo (MPRAGE) sequence with the following parameters: TR = 1,900 ms, TE = 2.52 ms, TI = 900 ms, flip angle = 90°, matrix size = 256 × 256, number of slices = 176, slice thickness = 1.0 mm, and voxel dimensions = 1 × 1 × 1 $mm^3$.

### 2. Stability of PMDF-TBA

To evaluate the stability of PMDF-TBA, we constructed multiple PMDF-TBAs using varying database sizes (ranging from 1 to 25 with a step size of 1) and assessed their prediction accuracy in estimating the sensitivity of brain regions for three widely used atlases (Brodmann, AAL2, and LPBA40) on unseen subjects. A leave-one-out cross-validation approach was employed for this purpose. In each cross-validation iteration, one subject was selected from the total of 48 subjects as the test subject, while the remaining 47 subjects constituted the candidate database. For each database size **m**, we randomly selected m subjects from the candidate database and calculated the corresponding PMDF-TBA(**m**). This PMDF-TBA(**m**) was then used to predict the sensitivity of the test subject, yielding a prediction error. The process was repeated 100 times, and the mean prediction error across these 100 tests was obtained for the given cross-validation iteration. After completing all 48 cross-validation iterations, we calculated the overall mean prediction error to indicate the prediction accuracy for PMDF-TBA(**m**). The results are demonstrated in Fig S6.

Table S1. Tissue optical properties

| | $\mu_a$(mm-1) | $\mu_s$(mm-1) | g | n |
|---|---|---|---|---|
| Scalp | 0.017275 | 0.72 | 0.01 | 1 |
| Skull | 0.011925 | 0.92 | 0.01 | 1 |
| CSF | 0.0025 | 0.01 | 0.01 | 1 |
| Gray matter | 0.0195 | 1.1 | 0.01 | 1 |
| White matter | 0.0169 | 1.35 | 0.01 | 1 |

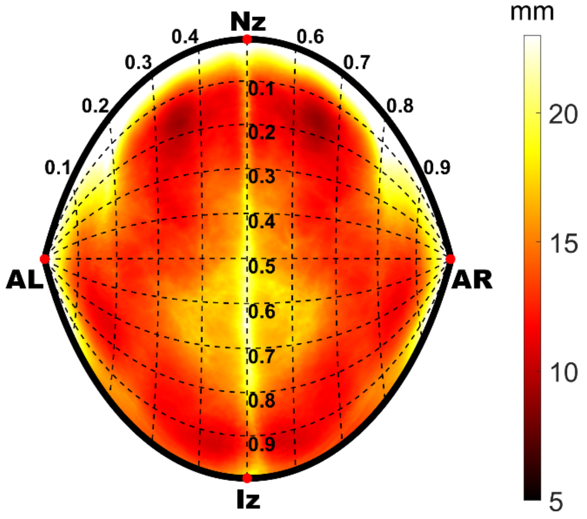


**Figure S1**. Group average scalp-to-cortex distance (in millimeters)

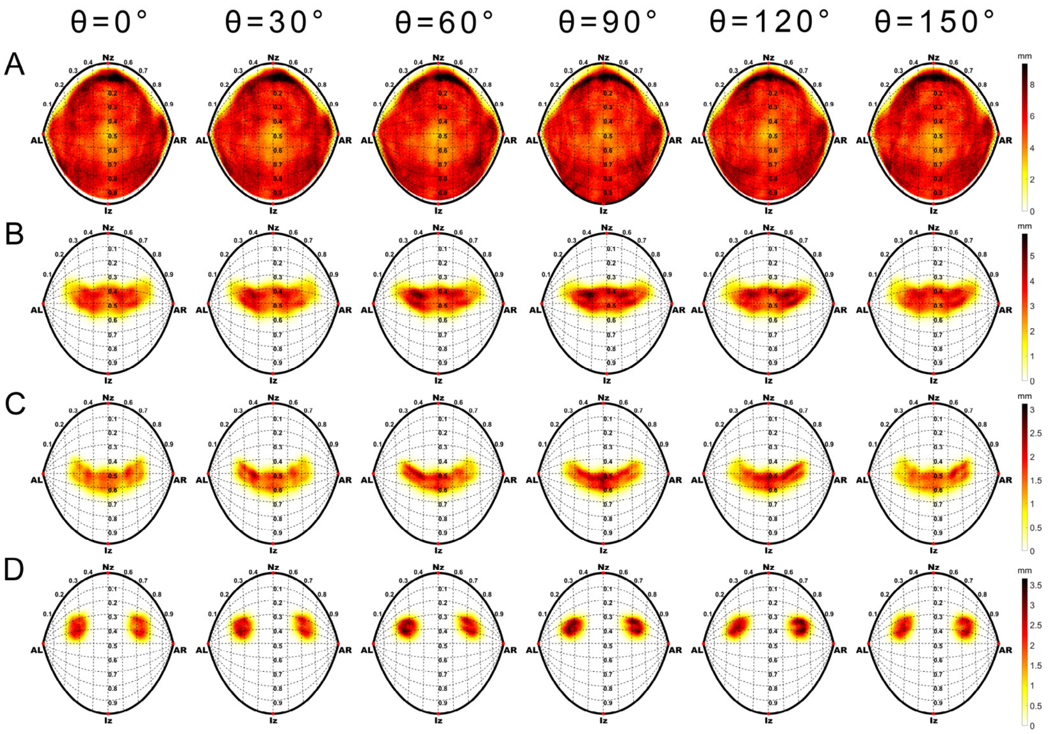


**Figure S2**. Standard deviation of sensitivity of gray matter (A), BA 6 (B), BA 4 (C) and BA 44 (D).

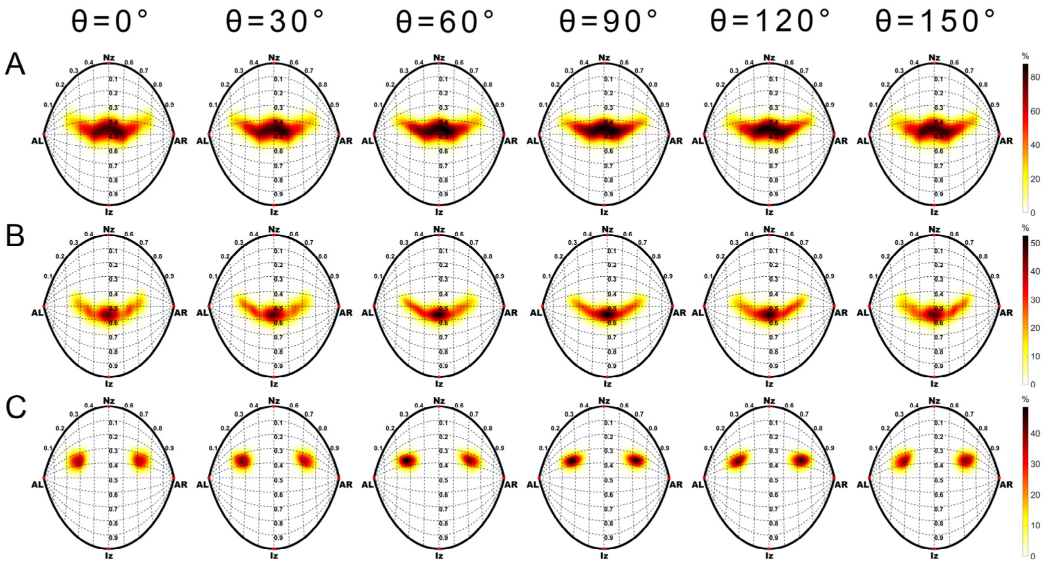


**Figure S3**. Specificity of BA 6 (A), BA 4 (B) and BA 44 (C).

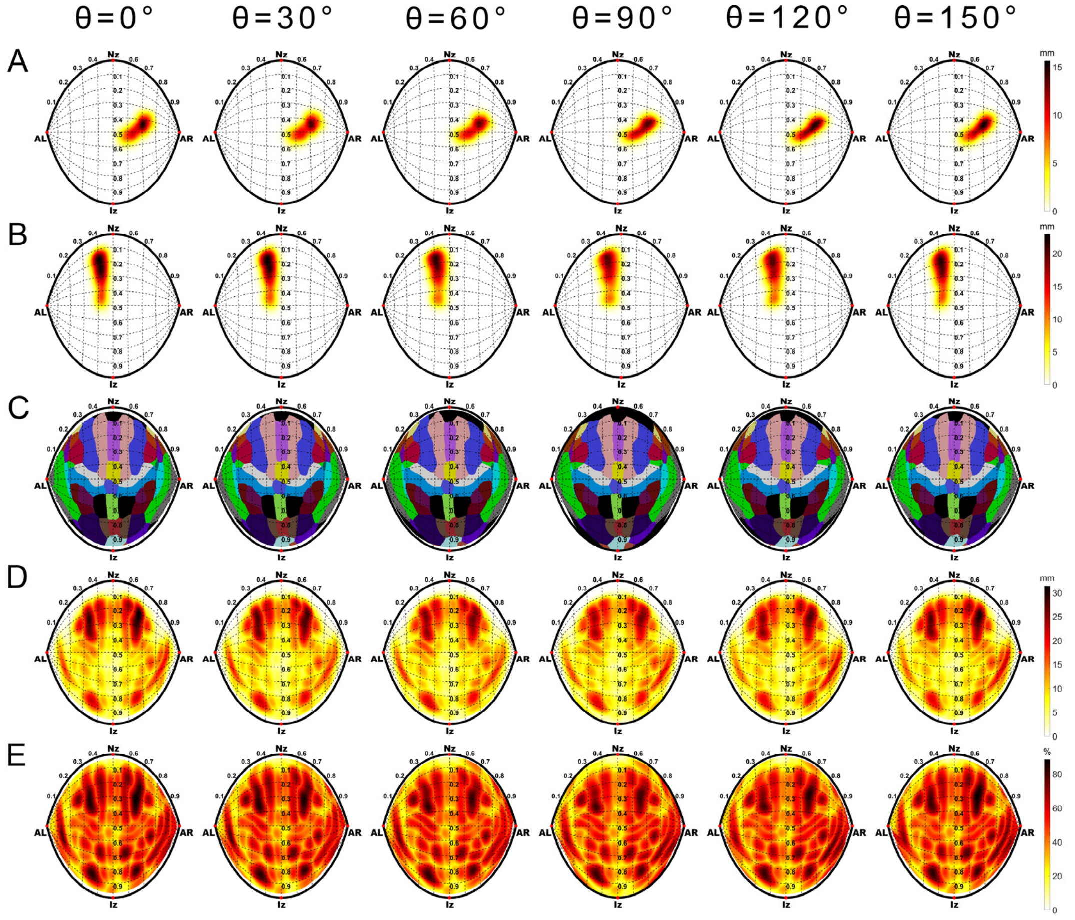


**Figure S4**. PMDF-TBA of AAL2. Sensitivity maps of right precentral gyrus(A), and left dorsolateral superior frontal gyrus (B). The maximum sensitivity label map (C) of the Brodmann areas, and the corresponding maximum sensitivity map (D) and maximum specificity map (E).

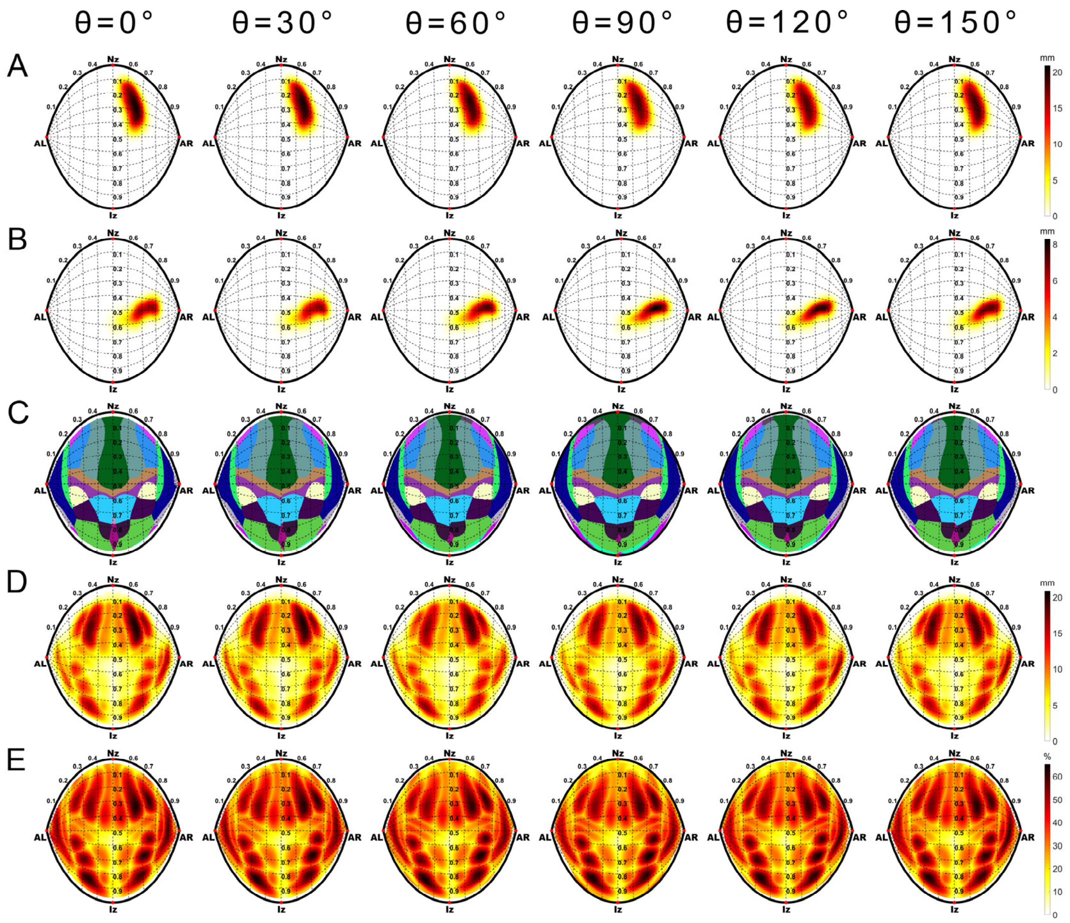


**Figure S5**. PMDF-TBA of LPBA40. Sensitivity maps of right middle frontal gyrus(A), and right postcentral gyrus (B). The maximum sensitivity label map (C) of the Brodmann areas, and the corresponding maximum sensitivity map (D) and maximum specificity map (E).

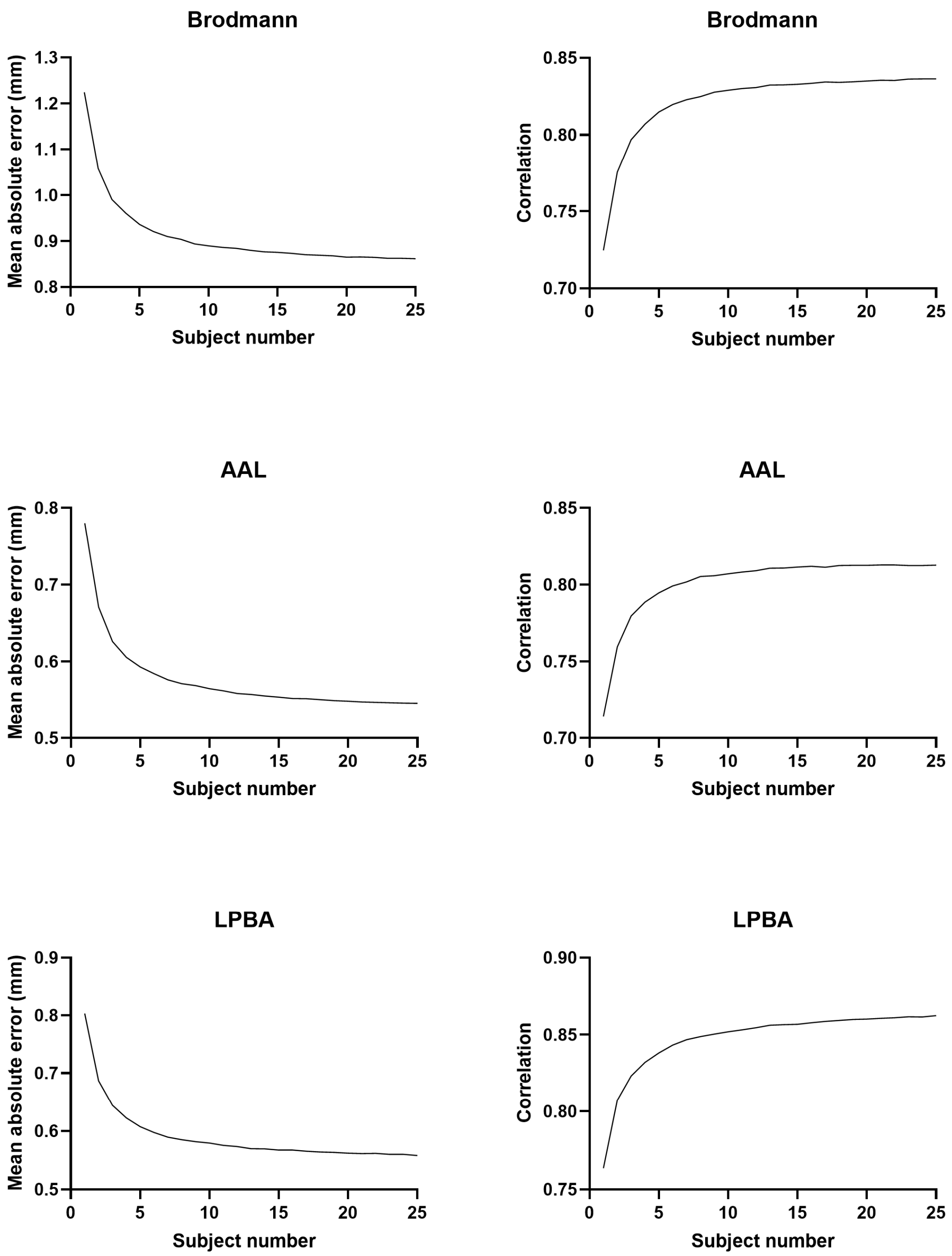


**Figure S6**. Relationship between database size and prediction error

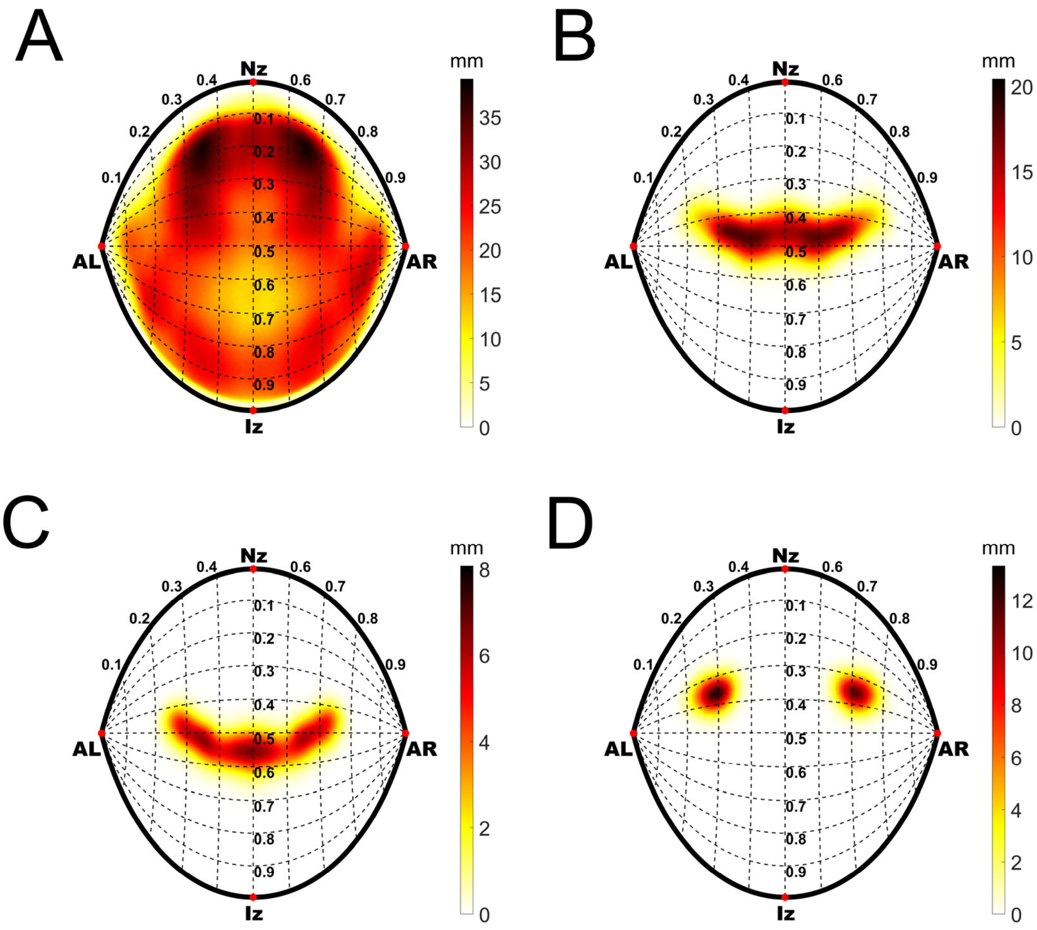


**Figure S7**. Orientation-averaged sensitivity maps

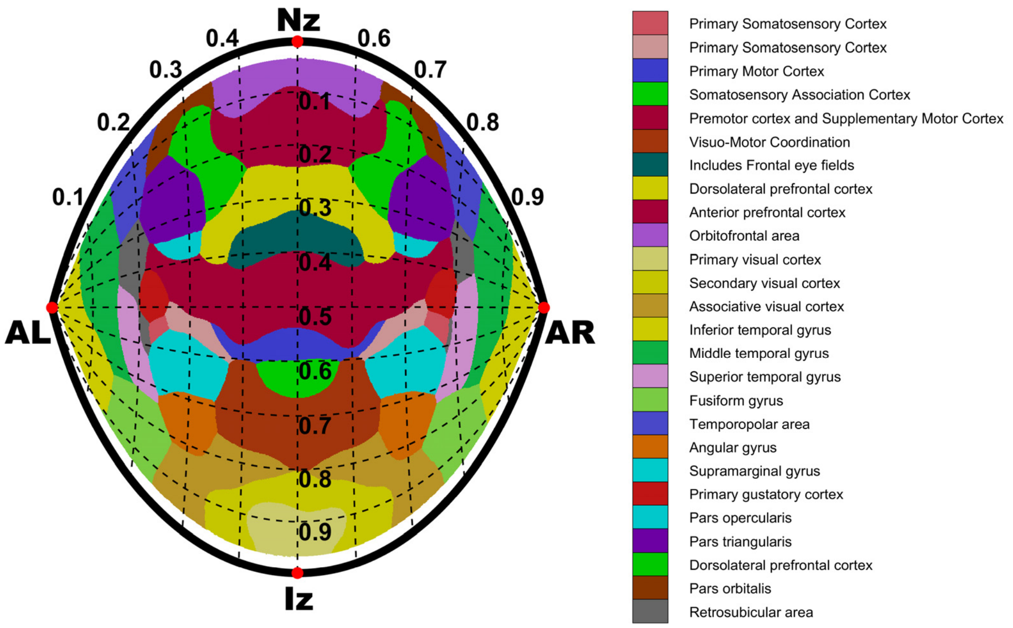


**Figure S8**. Names of maximum sensitivity labels (shown only for channels with θ = 0°).
